# *Tbx1* regulates extracellular matrix-cell interactions in the second heart field

**DOI:** 10.1101/267906

**Authors:** Daniela Alfano, Alessandra Altomonte, Claudio Cortes, Marchesa Bilio, Robert G. Kelly, Antonio Baldini

## Abstract

*Tbx1,* the major candidate gene for DiGeorge or 22q11.2 deletion syndrome, is required for efficient incorporation of cardiac progenitors (CPs) of the second heart field (SHF) into the heart. However, the mechanisms by which TBX1 regulates this process are still unclear. Here, we have used two independent models, mouse embryos and cultured cells, to define the role of TBX1 in establishing morphological and dynamic characteristics of SHF in the mouse. We found that loss of TBX1 impairs extra cellular matrix (ECM)-integrin-focal adhesion (FA) signaling in both models. Mosaic analysis in embryos showed that this function is non-cell autonomous and, in cultured cells, loss of TBX1 impairs cell migration and focal adhesions. Additionally, we found that ECM-mediated outside-in integrin signaling is disrupted upon loss of TBX1. Finally, we show that interfering with the ECM-integrin-FA axis between E8.5 and E9.5 in mouse embryos, corresponding to the time window within which TBX1 is required in the SHF, causes outflow tract dysmorphogenesis. Our results demonstrate that TBX1 is required to maintain the integrity of ECM-cell interactions in the SHF, and that this interaction is critical for cardiac outflow tract development. More broadly, our data identifies a novel TBX1 downstream pathway as an important player in SHF tissue architecture and cardiac morphogenesis.

## INTRODUCTION

The heart develops from populations of progenitor cells that are specified as early as gastrulation. *In vivo* labeling and genetic clonal analyses have identified at least two distinct populations, the first (FHF) and second (SHF) heart fields (1-5). These lineages derive from cells that independently activate *Mesp1* in the primitive streak and differentiate sequentially during cardiogenesis to give rise to distinct parts of the heart (6, 7). The SHF is part of a mesodermal population that gives rise to cardiomyocytes as well as other cell types, including branchiomeric skeletal muscles and some populations of endothelial cells. Overall, this multipotent cell population is referred to as the cardiopharyngeal mesoderm (CPM) lineage (8).

CPM/SHF progenitors migrate as part of the splanchnic pharyngeal mesoderm and undergo progressive differentiation to give rise to myocardium at the cardiac poles. Defective deployment of SHF cells results in a spectrum of common forms of congenital heart defects (CHD). The cellular and molecular genetic mechanisms by which progenitor cells are directed and sorted into the different cell types and regionalized in the pharyngeal apparatus towards the cardiac poles are only now beginning to emerge (9). Specifically, SHF progenitors destined to contribute to the arterial and venous poles of the heart are organized into an epithelial-like layer of cells, between embryonic day (E) 8 and E9.5, that forms the dorsal pericardial wall (DPW). The epithelial properties of the SHF are emerging as a critical regulatory step during the process of heart tube elongation, yet how regulators of SHF deployment impact on SHF cell biology remains poorly understood (10). *Tbx1* has been shown to be required for progenitor recruitment to the heart (11, 12) and its loss has been associated with disruption of epithelial polarity in the DPW (13), and reduced tension of the epithelial sheet (14), although the mechanisms by which these phenotypic anomalies occur remain unclear. Planar cell polarity (PCP) genes *Dvl2* and *Wnt5a* have been proposed to be part of this process by facilitating the incorporation, by intercalation, of mesenchymal cells into the epithelial sheet of the DPW (15-17). TBX1 regulates *Wnt5a* expression in the SHF (18), suggesting that at least some of the tissue architectural roles of TBX1 may operate through the non-canonical WNT/PCP pathways. However, *Wnt5a*^*-/-*^*;Tbx1*^*-/-*^embryos have much more severe SHF-derived heart defects than the individual mutants, providing genetic evidence that the two genes also have non-overlapping functions in the SHF (18).

Here, we use two independent models, mouse embryos and cultured cells, to show that *Tbx1* regulates the extra cellular matrix-integrin-focal adhesion axis and alters the properties of splanchnic mesoderm, both in epithelial cells of the DPW and in the adjacent mesenchyme. We demonstrate that the epithelial layer of the SHF has an apical adhesion domain that includes immunoreactivity for E-cadherin, beta-catenin, paxillin, and actomyosin, and this domain is disrupted in *Tbx1*^-/-^embryos. Moreover, our results indicate that TBX1 is required to sustain ECM-intracellular signaling and that integrin-focal adhesion integrity is critical for cardiac outflow tract development. These findings identify TBX1 regulation of the ECM-focal adhesion axis as a critical component of early heart morphogenesis and candidate pathway in the etiology of congenital heart defects.

## RESULTS

### The ECM-Integrin-FA axis is altered in the SHF of *Tbx1* mutants

Despite evidence for the importance of the epithelial features of SHF cells in the DPW, how these properties are regulated is unclear. In particular how the ECM and cell adhesion are coordinated during SHF deployment has not been studied. We set out to test a group of markers critical for cell dynamics in the SpM of WT and *Tbx1* mutant embryos; specifically, we investigated markers of the extracellular matrix (ECM), integrin, and focal adhesion. At E9.5 COLI, the most abundant fibrillar collagen (encoded by the *Col1a1* gene), was expressed in the SpM, laterally and posteriorly to the pericardial cavity (Fig. 1A). COL1 also accumulates in the mesenchymal tissue adjacent and abutting to the basal side of the epithelial-like layer of the SHF (eSHF), where it formed a rudimentary and discontinuous basal lamina, but was not observed laterally or apically to the eSHF cells (Fig. 1C). In *Tbx1*^-/-^ embryos, COLI immunostaining was increased in the SpM and we noted a broader and mislocalized distribution in the eSHF where the collagen fibrils were randomly distributed around the cells, rather than being restricted to a basal lamina (Fig. 1B, D and Supplementary Fig. 1-2).

**Figure 1.**
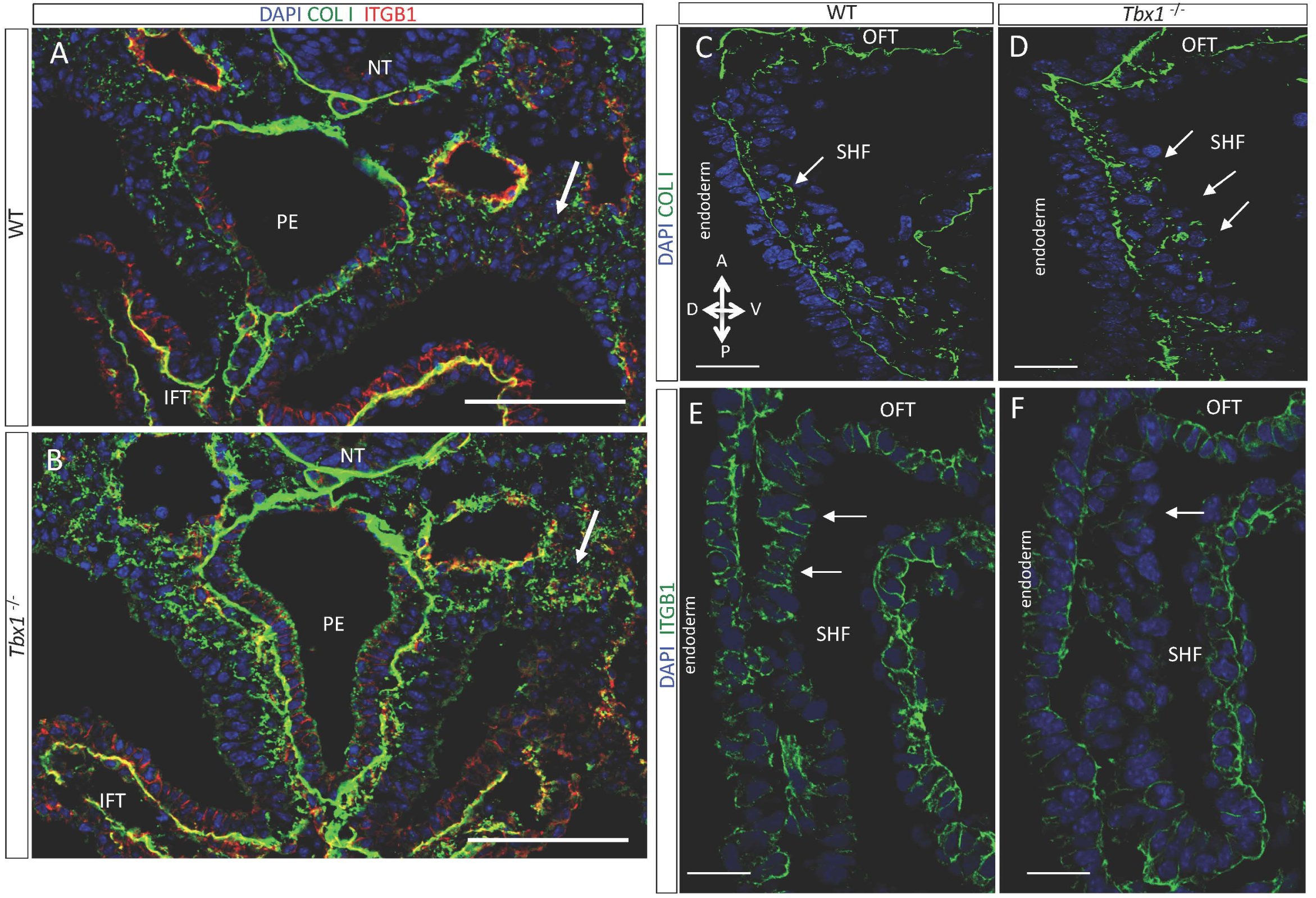
Collagen I and integrin β1 are altered in *Tbx1* mutants. **(A-B)** Transverse sections of E9.5 embryos showing the distribution of COLI in the SpM. Arrows indicate COLI accumulation in the *Tbx1*^*-/-*^ mutant (scale bar, 100*μ*m). **(C-D)** Confocal images of COLI immunostaining at SHF level of WT (C) and *Tbx1*^-/-^ embryos (D) (sagittal sections, E9.5). In the WT, COLI forms a rudimentary and discontinuous basal lamina while in *Tbx1*^-/-^ embryos its distribution is abnormal, showing an accumulation between eSHF cells (see arrows) (scale bar, 40*μ*m). (**E-F**) ITGB1 immunostaining of E9.5 WT and *Tbx1*^-/-^ embryos. In the WT embryo, the signal is present in the anterior SHF and is not detectable in *Tbx1*^-/-^ embryo (see arrows) (scale bar, 20*μ*m). NT, neural tube; IFT, inflow tract; PE, pharyngeal endoderm; OFT, outflow tract; A, anterior; P, posterior; V, ventral; D, dorsal.

ITGB1, the *β*1 integrin subunit, was expressed in the anterior SHF (here defined as the anterior half of the DPW, close to the arterial pole of the heart), but not in the posterior SHF (Fig. 1E). ITGB1 was principally localized at cell-cell contacts and at the basal side of eSHF cells (Fig. 1E and Supplementary Fig. 2), and to a lesser extent at the apical region. ITGB1 was decreased in *Tbx1*^-/-^ embryos compared to WT embryos (Fig. 1F and Supplementary Fig. 2).

**Figure 2.**
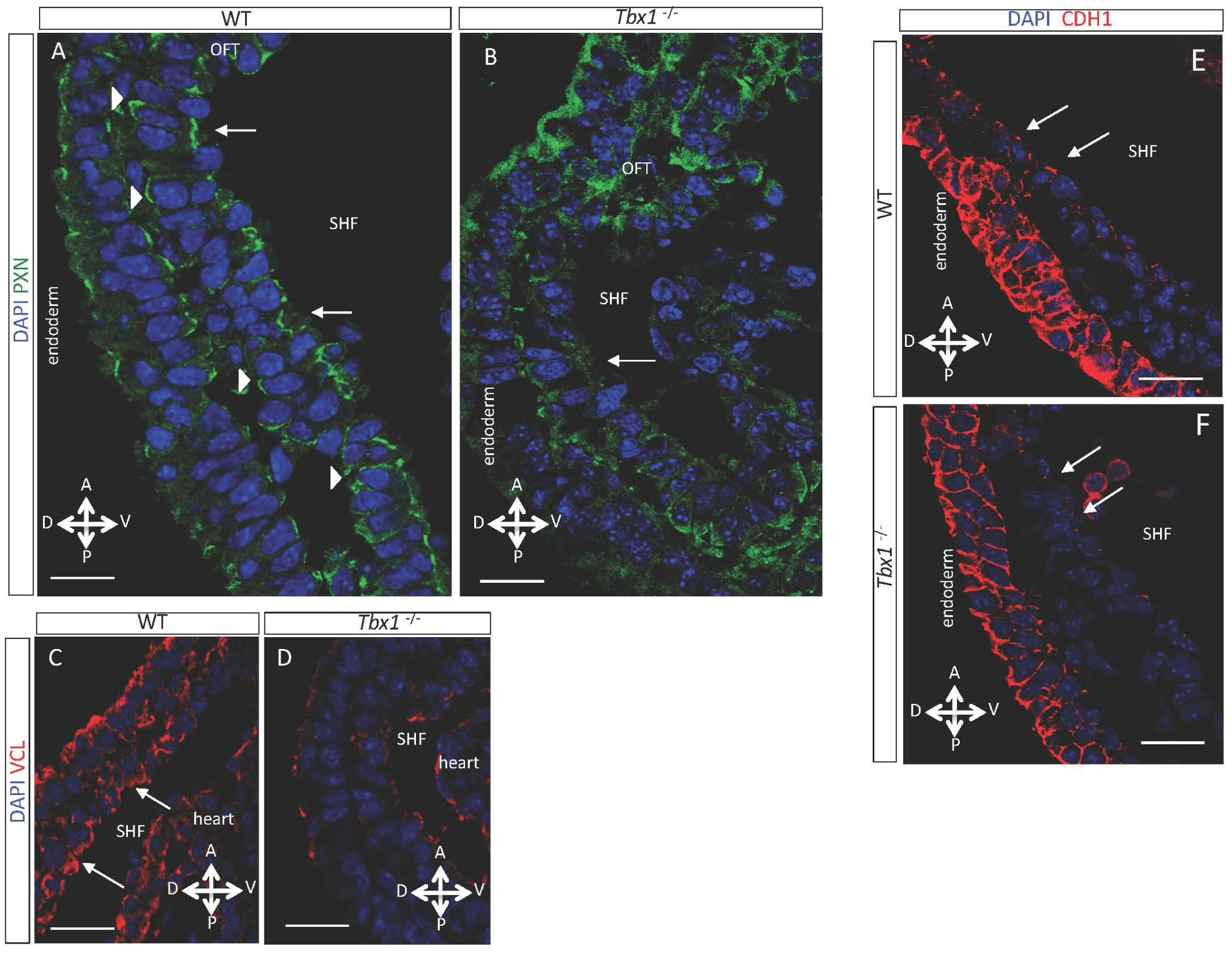
TBX1 regulates FA and cell-adhesion proteins at the eSHF. **(A-B)** Immunofluorescence on sagittal sections of E9.5 wild-type (WT) and *Tbx1*^-/-^ embryos, showing the distribution of PXN. (A) In the WT embryo PXN accumulates predominantly at the apical surface and cell junctions (arrows), forming a nearly continuous line in the apical membrane of eSHF cells. PXN staining was also found in the basal domain (arrowheads) of eSHF cells. In the endoderm PXN staining marks both apical and basal domains. (B) In the *Tbx1*^-/-^ embryo, the PXN pattern in the eSHF cells is completely abolished (scale bar, 20*μ*m). **(C-D)** Immunostaining of VCL on sagittal sections of an E9.5 WT and *Tbx1*^-/-^ embryos. (C) In the WT embryo, VCL displays an apical domain in the eSHF cells (arrows), similarly to the PXN domain. (D) In the *Tbx1*^-/-^ embryo, this characteristic pattern of VCL is severely affected and intensity of staining is reduced (scale bar, 30*μ*m). (**E-F**) In a WT embryo, CDH1 is observed on the apical-lateral domains of the eSHF monolayer, predominantly in the anterior region of the SHF (arrows). In the *Tbx1*^-/-^ embryo, the CDH1 signal is reduced or nearly abolished in the posterior-lateral regions of SHF (scale bar, 15*μ*m). OFT, outflow tract; A, anterior; P, posterior; V, ventral; D, dorsal.

Considering that transduction of integrin signaling is associated with focal adhesions, we used IF with an antibody for the FA marker Paxillin (PXN) and found that it is expressed in the SHF and is predominantly localized in the apical and, to a lesser extent, the basal side of cells in the eSHF (Fig. 2A). A similar apical and basal distribution of PXN was also visible in other epithelia, such as the pharyngeal endoderm (Fig. 2A). In the eSHF, we found that apical PXN staining partially overlapped with cell-cell adhesion complex proteins E-cadherin (CDH1) and beta-catenin (CTNNB1) (Fig. 3A-H and Supplementary Movie 1, Supplementary Movie 2). Specifically, the immunostaining overlap was observed in the apical-lateral region of eSHF cells, at the level of cell-cell junctions (Fig. 3A-H), where tight and adherens junction proteins accumulate. In the eSHF we observed a similar pattern with another FA protein, Vinculin (VCL, Fig. 2C). PXN expression in *Tbx1*^-/-^ embryos was reduced and, strikingly, its apical-lateral distribution could not be detected (Fig. 2B). Likewise, VCL was also reduced (Fig. 2D). Moreover, CDH1 was reduced in *Tbx1*^*-/-*^ mutants, especially in posterior-lateral regions of the DPW (Fig. 2E-F). Cortical actin staining was also reduced in *Tbx1*^*-/-*^ mutants (Supplementary Fig. 3A-B), suggesting impaired assembly of junctional F-actin networks. The actin-motor non-muscle myosin IIB (NMIIB, a.k.a. MYH10) regulates both actin rearrangements and FAs formation and is required for OFT morphogenesis (19-21). We found that NMIIB predominantly decorates the apical side and, to a lesser extent, the lateral side of eSHF cells, while in *Tbx1*^-/-^ embryos, this specific localization was lost and the protein was randomly distributed around the eSHF cells (Supplementary Fig. 3C-D). The altered distribution of NMIIB reinforces the idea that TBX1 supports the ability of the eSHF to undergo morphogenetic changes, acquiring epithelial features which are essential for proper heart morphogenesis (13, 14).

**Figure 3.**
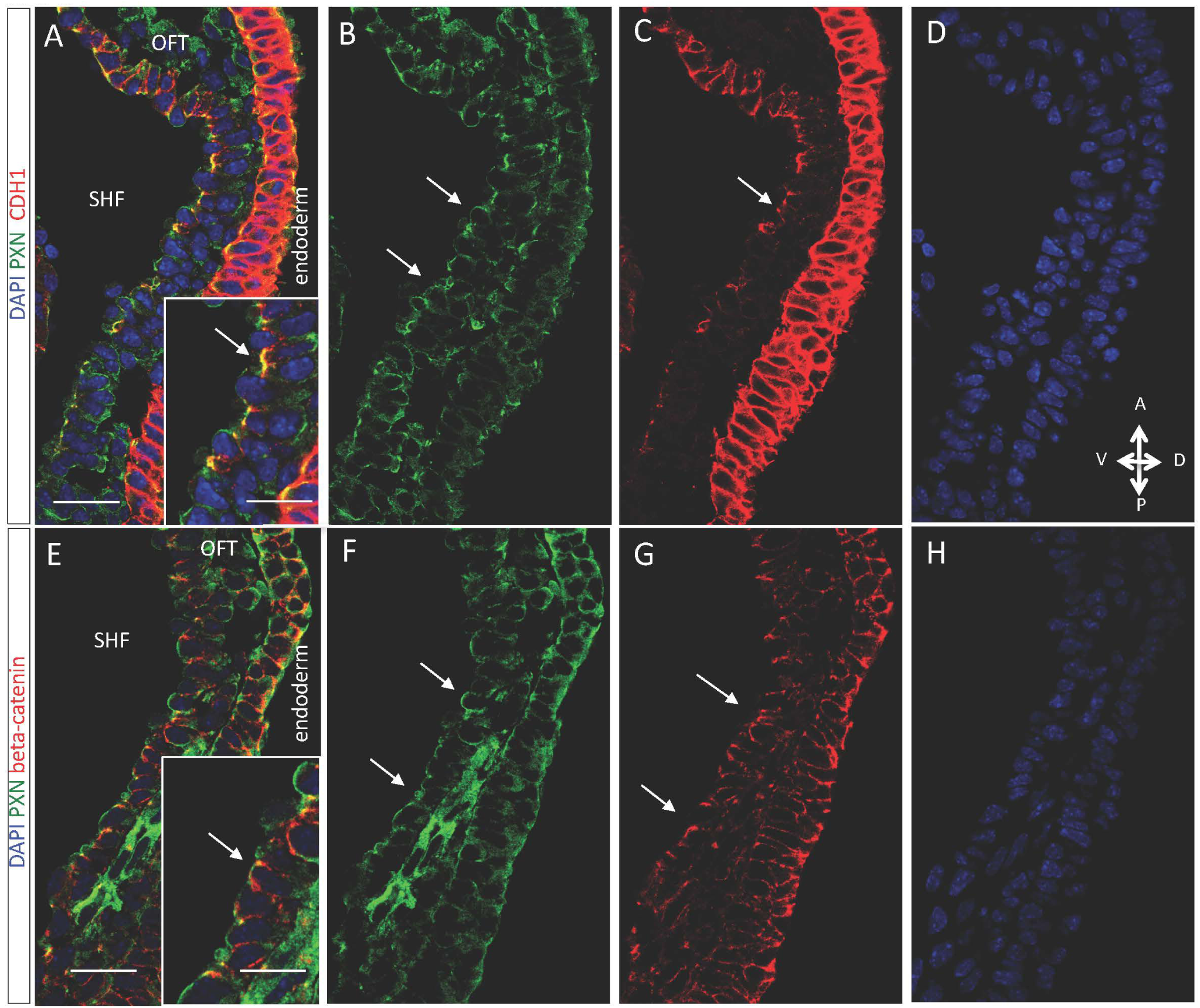
Apical-junctional co-localization of PXN and CDH1 in the eSHF. (**A-D**) Immunofluorescence of sagittal sections of E9.5 WT embryos showing the distribution of PXN and the cell-cell junction proteins CDH1 and beta-catenin. PXN accumulates in the apical-lateral domain of eSHF cells and partially overlaps with CDH1-positive junctions (arrow) or (**E-H**) with beta-catenin-positive junctions (arrows) (scale bars, 40*μ*m). High-magnification views are boxed in A and E (scale bar, 10*μ*m). OFT, outflow tract; A, anterior; P, posterior; V, ventral; D, dorsal.

ECM- and cell-cell interactions are critical for cell orientation and apico-basal polarization. Considering that cellular elements like the centrosome or the Golgi apparatus are polarised in epithelial cells, we used Giantin, a Golgi marker, to assay apicobasal polarity of eSHF cells. In WT embryos, the Golgi apparatus of most eSHF cells (∼80%) is apically positioned (Supplementary Fig. 3E-G), whereas in *Tbx1*^-/-^ embryos approximately only ∼60% of cells presented an apical localisation (Supplementary Fig. 3E-G). This polarization defect was observed predominantly in the anterior eSHF (Supplementary Fig. 3G). Overall, these findings identify novel features of the eSHF and *Tbx1* null epithelial phenotype and show that loss of function of *Tbx1* causes a multifaceted cell phenotype related to impaired ECM-cell interactions.

### Cell non-autonomous functions of TBX1 in the SHF

We reasoned that if loss of *Tbx1* alters the overall tissue organization by altering ECM-cell interactions, the deletion of *Tbx1* in a small number of cells within the tissue should be insufficient to alter the tissue architecture. To test this, we used E9.5 *Tbx1*^*mcm*/flox^;*R26R*^mT-mG^ embryos in which Tamoxifen (TM) induction had activated Cre recombination at E7.5. In these embryos GFP+ cells are functionally *Tbx1* null, since the *Tbx1*^*mcm*^ allele is a null allele (11). Because of the low efficiency of this MerCreMer driver, GFP+ cells are relatively rare. We used *Tbx1*^*mcm*/+^;*R26R*^mT-mG^ (*Tbx1* heterozygous) embryos as controls. Results showed that NMIIB staining of GFP+ cells in *Tbx1*^*mcm*/flox^;*R26R*^mT-mG^ embryos was indistinguishable from that of the surrounding cells of the same embryos or control embryos (Supplementary Fig. 4A-B and Supplementary Fig. 4E). Furthermore, CDH1 distribution appeared homogeneous between GFP+ and surrounding cells (Supplementary Fig. 4C-D and Supplementary Fig. 4F). These findings indicate that the anomalous distribution of NMIIB or the alteration of CDH1 levels are not cell-autonomous phenotypes, suggesting that they are secondary to ECM disorganization and/or that there is a rescue-by-neighbors effect.

**Figure 4.**
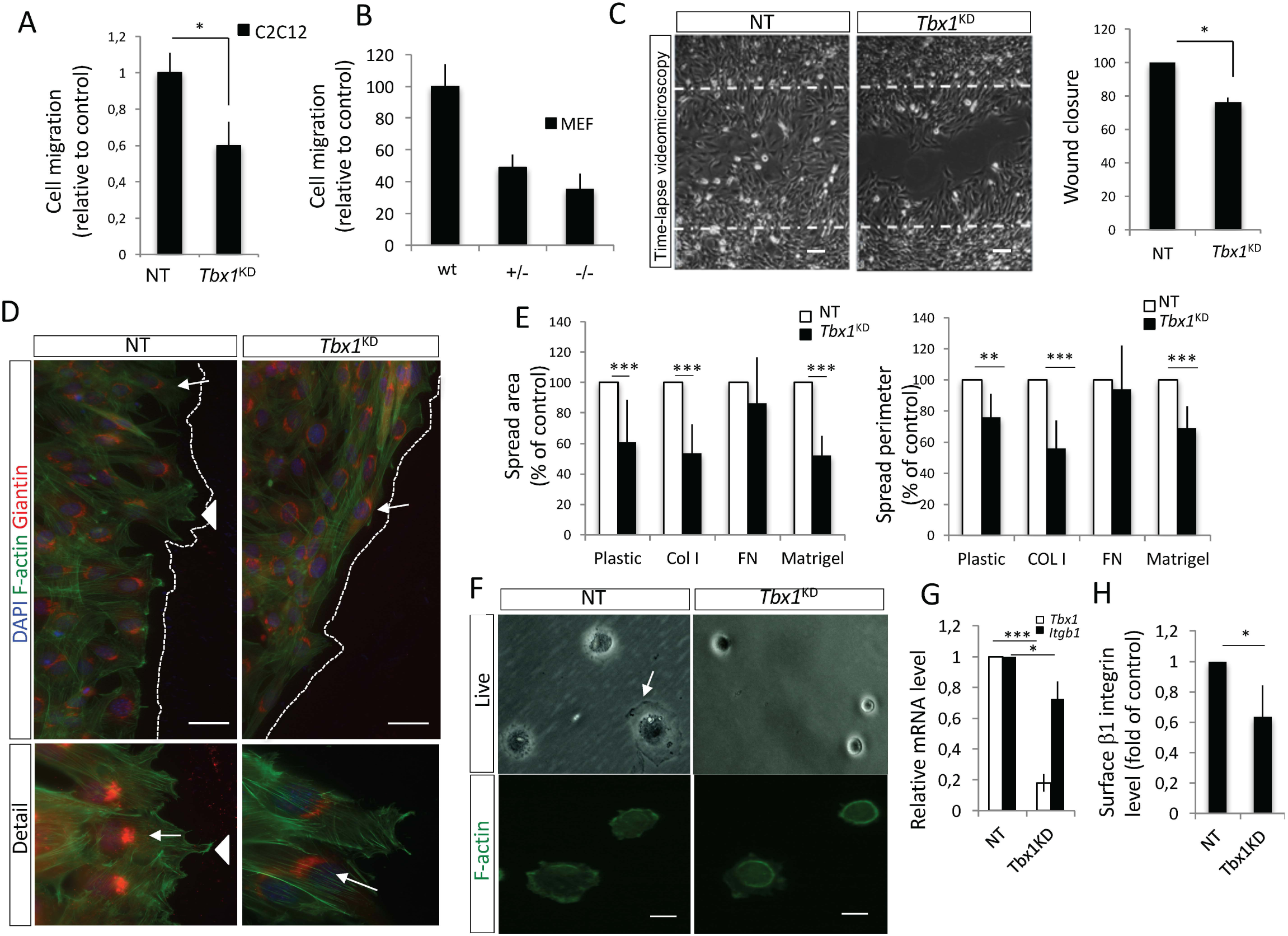
TBX1 regulates cell migration and polarity. **(A-B)** siRNAs-transfected C2C12 cells (A) or MEF (B) were plated on COLI-coated Transwell filters containing 10% FBS in the bottom well. The migration values are the means ± s.d. of three experiments performed in triplicate, normalized to the number of migrating cells in the absence of FBS, shown as fold change relative to control (\**p*< 0.05). **(C)** Left panel: phase contrast images taken from a time-lapse series at 16 hours after scratch wounding of control (NT) and *Tbx1*^*KD*^ cells (scale bar: 100 µm); right panel: Scratch wound area, showed as percentage of the initial wound area as determined using ImageJ software. The values are the means ± s.d. of three independent experiments (\**p*< 0.05). **(D)** Images of immunofluorescence with anti-Giantin and staining with phalloidin (F-actin) (upper panels) and respective high magnification views (lower panels); arrowheads indicate lamellipodia structures. Cells were transfected with the indicated siRNA, and after 42 hrs, a wound was made using a pipette tip and cells were allowed to migrate and polarize for approximately 5 hrs, then they were fixed and processed for immunofluorescence; arrows indicate the Giantin signal (scale bar, 50*μ*m). (**E**) Quantification of cell area (left panel) and cell perimeter (right panel) of siRNAs-transfected C2C12 cells after seeding on plastic uncoated or coated with COLI or FN for 30 min (mean ± s.d. from four different experiments with >300 cells per condition). NT: non-targeting control siRNA. Cell area and perimeter were determined by using ImageJ software. Values are shown relative to NT (\*\*\**p* < 0.001; ** *p* < 0.01). **(F)** Representative phase-contrast images of cells plated on COLI at sparse density for 20 min (top panels) or Phalloidin staining of spread cells after 30 min on COLI (bottom panels); white arrow represents lamella (scale bar, 20*μ*m). **(G)** After 48hrs from siRNAs transfection, C2C12 cells were subjected to RNA extraction for qPCR. Relative RNA levels of ITGB1 were normalized to Rpl13a expression. Asterisks above bars indicate significant differences compared with controls (\**p*<0.05). Error bars: s.d.; *n*=3. **(H)** C2C12 cells were transfected with siRNAs, and stained for surface levels of ITGB1, then analyzed by flow cytometry. Values are the mean fluorescence of the population ± s.d. of two separate experiments each performed in duplicate.

### *Tbx1* regulates cell migration, polarization, and adhesion in cultured cells

To address the role of TBX1 on cell dynamics, we decided to use cell culture models that allow for manipulation of the FA-integrin pathway. To understand whether TBX1 is involved in cell movements, we first performed a generic cell migration assay in a chemotactic gradient of serum. For this, we used two independent cell lines that express *Tbx1*: C2C12 undifferentiated myoblast cells and primary mouse embryonic fibroblasts (MEFs). For C2C12 cells, we used RNA interference to knock down (KD) *Tbx1* expression, while MEFs were obtained from WT, *Tbx1*^+/-^ and *Tbx1*^-/-^ embryos. Results showed that cell migration was significantly reduced following either TBX1 dosage reduction or KD (Fig. 4A-B and Supplementary Fig. 5A). We also used a two dimensional assay based on closure of mechanically inflicted wounds in confluent cell monolayers, monitored by time-lapse microscopy for 24 hrs. *Tbx1* depletion reduced C2C12 cell migration into the wound area (Fig. 4C); this effect was not due to differences in cell proliferation because we did not find a significant change in cell proliferation in the time window within which the migration assay was performed (Supplementary Fig. 5B). Furthermore, we found that migrating *Tbx1*-depleted (*Tbx1*^*KD*^) cells plated on COLI failed to polarize properly, as they exhibited aberrant repositioning of the Golgi apparatus (Fig. 4D), consistent with *in vivo* data. The number of polarized cells was reduced by 80% in *Tbx1*^*KD*^ cells compared with control cells (Supplementary Fig. 5C). Phalloidin staining revealed that control cells formed lamellipodia at the edge of the wound but *Tbx1*^*KD*^ cells showed fewer lamellipodial protrusions (Fig. 4D and Supplementary Fig.6A-E), indicating that TBX1 is required for the formation of these structures, which produce the driving force for migration. Moreover, IF analysis showed that in control cells alpha-tubulin is enriched at the leading edge, while in *Tbx1*^*KD*^ cells the microtubule distribution is more uniform around the cells (Supplementary Fig. 5D-E and Supplementary Fig. 6B-E confirming a polarization defect.

**Figure 5.**
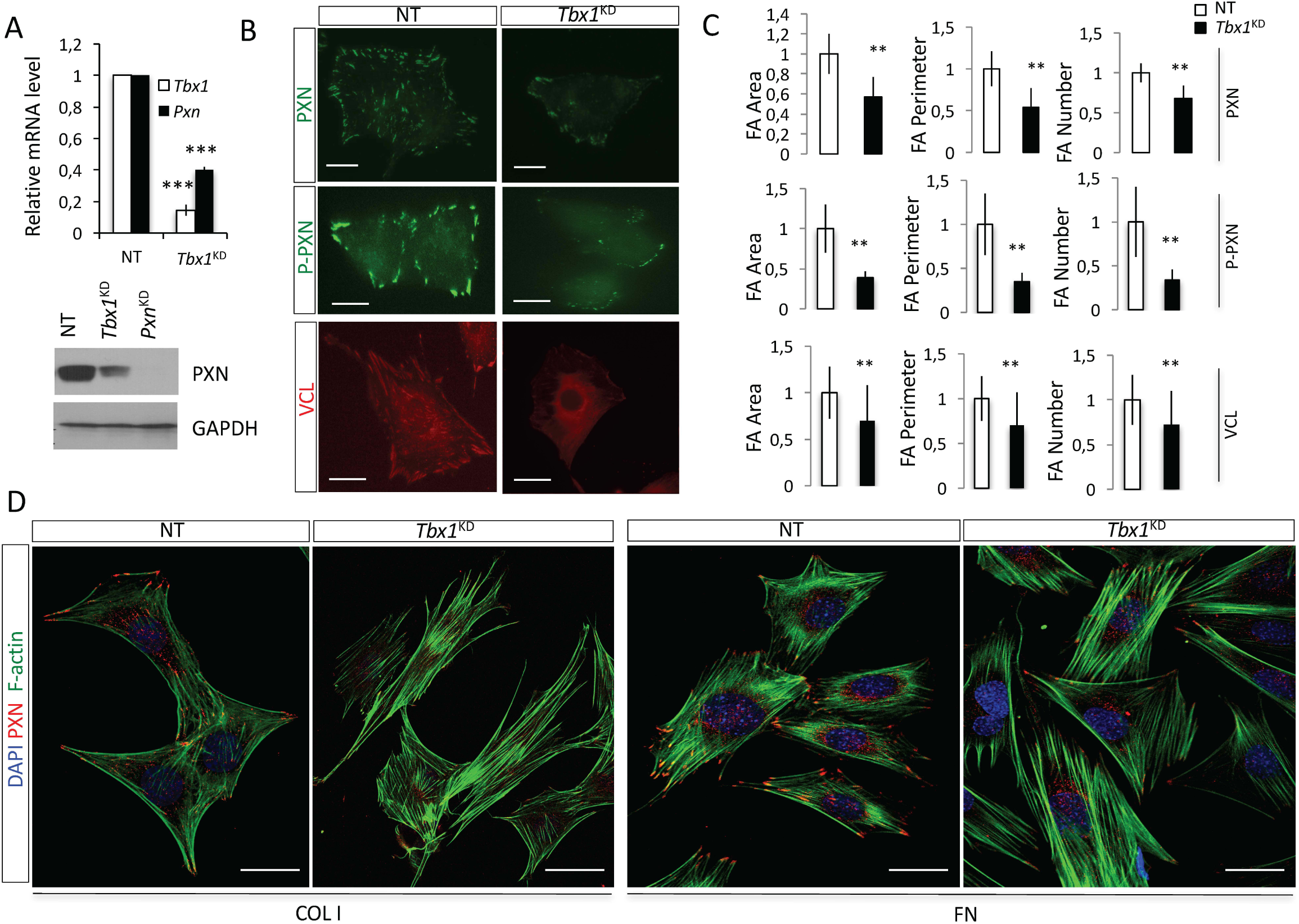
TBX1 controls FA formation in C2C12 cells. **(A)** After 48hrs from siRNAs transfection, C2C12 cells were subjected to RNA and protein extraction for qPCR (top panel) and western blot (bottom panel) assays, respectively. Top panel: relative RNA levels were normalized to Rpl13a expression. Asterisks above bars indicate significant differences compared with controls (\*\*\**p*<0.001). Error bars: s.d.; *n*=3. Bottom panel: immunostaining of PXN and GAPDH (loading control). Data are representative of three independent experiments. **(B)** Representative pictures of cells immunostained with PXN (top panels), phosphorylated PXN (P-Pxn Y118) (middle panels) or VCL antibodies (bottom panels). (**C**) Graphs show the quantification of area occupied by FAs, perimeter covered by FA and the number FAs/per cell, relative to the total corresponding spreading values (NT, white columns; *Tbx1*^*KD*^, black columns). All quantitative analyses were performed separately, considering FAs positive for anti-PXN (>100 cells), anti-P-PXN (>40 cells) or anti-Vcl (>60 cells). The quantitative analysis was carried out by Cell Profiler software (mean ± s.d. from three different experiments) (scale bars, 25*μ*m). Values are shown relative to NT control cells (\**p* < 0.05; \*\**p* < 0.01). **(D)** Confocal images of immunofluorescence staining of cells seeded on ECM proteins showing FAs and focal complexes. Nuclei were stained with DAPI. (scale bars: 25*μ*m).

**Figure 6.**
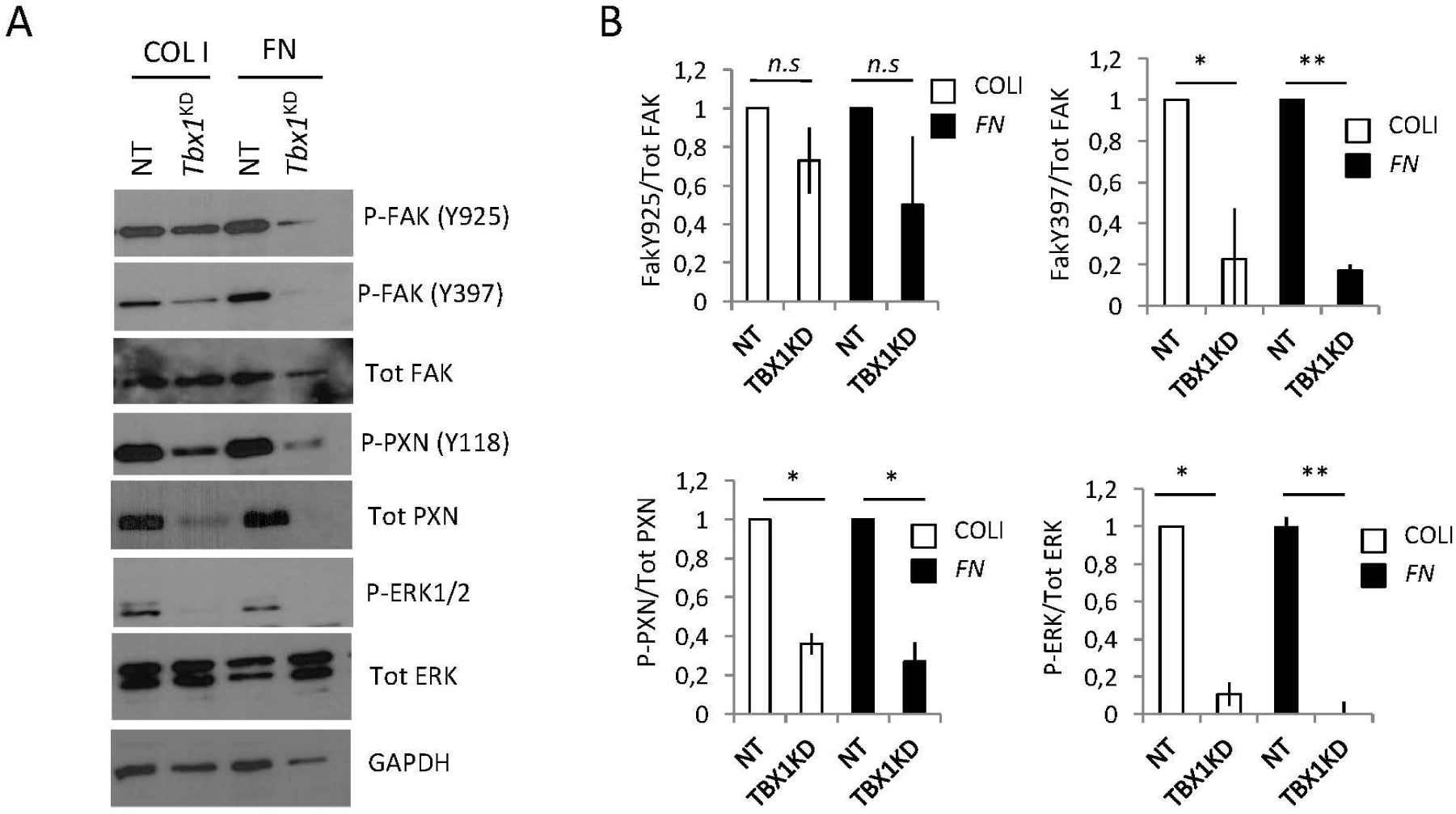
*Tbx1*-depleted cells fail to respond to ECM-FAs signalling. **(A)** After 48hrs from siRNAs transfection, C2C12 cells were plated on COLI or FN and lysed after 20 min. Cell lysates were analyzed by immunoblotting with antibodies to the indicated phosphorylated and to total protein levels of the respective protein. P-PXN, P-FAK and P-ERK1/2 levels were normalized to total PXN, FAK and ERK1/2 levels, respectively. GAPDH was used as a loading control. (**B**) Histograms show densitometric analysis of immunoblots. Values are levels of the normalized phosphorylated proteins relative to those in control cells (NT) (*n*=3 for each protein; means ± s.d. \**p*<0.05, ** *p*<0.01 versus control).

The migration impairment might be due to defective interactions with the substrate (22). The cell spreading assay in tissue culture is useful to test cell adhesion defects and, in particular, defects regarding the formation and disassembly of initial adhesive cell contacts. C2C12 cells were plated on a variety of substrates, specifically COLI, Fibronectin (FN), Matrigel, or uncoated plastic. We then evaluated cell spread area after 20 min in culture. We found that *Tbx1*^*KD*^ C2C12 cells had a significantly reduced spread area and perimeter compared to non-target (NT) siRNA-treated control cells on all substrates, except FN (Fig. 4E). *Tbx1*^*KD*^ cells appeared rounded and less spread compared to control cells (Fig. 4F). *Tbx1* depletion did not alter significantly the size of cells in suspension (Supplementary Fig. 5F). Furthermore, we found that, consistent with in vivo data, the expression of integrin *Itgb1* was decreased in *Tbx1*^*KD*^ cells (Fig. 4G) and ITGB1 was reduced in both the membrane surface and in FAs, as shown by flow cytometry and IF, respectively (Fig. 4H and Supplementary Fig. 5G-H). In addition, *Tbx1*^*KD*^ cells showed a significant increase of aberrant protrusions (Supplementary Fig. 5I-M), a phenotype previously shown to be associated with loss of Focal Adhesion Kinase (FAK) or PXN (23, 24). Indeed, PXN, which is a major adapter protein of FAs, was down regulated at both mRNA and protein levels in *Tbx1*^*KD*^ cells (Fig. 5A). Given these results, we tested for possible FA abnormalities. We analyzed control and *Tbx1*^*KD*^ cells in two different conditions of spreading that stimulate focal complex and adhesion formation (here collectively referred to as FAs): 1) in acute serum-induced spreading, or 2) by plating cells on different ECM compositions (COLI or FN). Results, obtained by PXN or phospho-PXN immunostaining, revealed that *Tbx1*^*KD*^ cells had a reduced number of FAs/cell compared to control cells, in both conditions (Fig. 5B-C, upper and middle panels and Fig. 5D). Moreover, in *Tbx1*^*KD*^ cells, the area and perimeter occupied by FAs relative to the total spread area and perimeter of the cell was significantly lower compared to control cells (Fig. 5B-C upper and middle panels). Similarly, VCL, which is a membrane-cytoskeletal protein in FA plaques, highlighted elongated fibrillar adhesions over the entire ventral surface of control cells by IF, whereas in *Tbx1*^*KD*^ cells, the VCL staining was predominantly restricted to shorter plaques and we rarely observed elongated FAs (Fig. 5B, lower panels). Quantitative analysis of VCL IF confirmed that the number and size of FAs were reduced in *Tbx1*^*KD*^ compared to control cells (Fig. 5C, lower panels).

Overall these findings reveal that TBX1 regulates ECM-integrin-FA signaling and that loss of function of TBX1 has major functional consequences for cell migration and spreading.

### TBX1 is a positive regulator of ECM-mediated signaling

The concomitant alteration of ECM, integrin, and FA protein expression and/or distribution in *Tbx1* mutant embryos suggests that TBX1 may regulate ECM-cell signaling. Integrins can signal through the cell membrane in either direction: the extracellular binding activity of integrins is regulated from the inside of the cell (inside-out signalling), while the binding of the ECM elicits signals that are transmitted into the cell (outside-in signalling) (25, 26). To determine the consequences of *Tbx1* silencing on outside-in signalling, we analyzed the activation of integrin-stimulated signalling cascades in C2C12 cells following adhesion to ECM proteins COLI and FN. In particular, we tested the principal components of PXN-mediated ECM-dependent signalling: FAK, extracellular-signal regulated kinases (ERK1/2), and Cofilin1 (CFL1). CFL1 is a critical cortical actin regulator, which severs actin filaments to increase the number of sites for actin polymerization in lamellipodia and is inhibited by adhesion signalling (including PXN-signalling) through phosphorylation of Ser3 (27). ECM engagement induced phosphorylation of FAK, ERK1/2, and CFL1 (Fig. 6A-B and Supplementary Fig. 6F) in control cells plated on COLI or FN for 30 min. In *Tbx1*^*KD*^ cells, the phosphorylation of PXN (Tyr118), FAK (Tyr397 and Tyr925), CFL1 (Ser3) and ERK1/2 were all strongly reduced (Fig. 6A-B). Consistent with this result, P-ERK expression has been shown to be reduced in the eSHF of *Tbx1*^-/-^ embryos (15). The defect of signalling cascade activation was only evident in ECM-plated *Tbx1*^*KD*^ cells (Supplementary Fig. 6G). These results indicate that loss of TBX1 affects at least two different aspects of ECM signalling: actin dynamics (CFL1) and cell adhesion (FAK, PXN, ERK).

### The integrity of the ECM-integrin-FA axis is required for OFT morphogenesis

To test whether interference with the ECM-integrin-FA axis between E8.5 and E9.5 (the critical time window for TBX1 function in the SHF) has consequences for OFT morphogenesis, we used chemical inhibitors in an embryo culture model (28). To this end, we used the disintegrin Echistatin (a potent integrin inhibitor), and a more specific inhibitor 6-B345TTQ, which blocks the alpha4 integrin-PXN interaction (29, 30). E8.5 WT embryos were cultured for 20 hours in the presence of inhibitor or vehicle (control) and OFT length and morphology were evaluated. Both inhibitors caused a significant reduction of OFT length compared to control embryos (Fig. 7G and Supplementary Fig. 7C). Additionally, histological analyses and N-Cadherin immunofluorescence of cultured embryos indicated that despite OFT dysmorphogenesis, epithelial integrity was not compromised (Fig. 7D-F).

**Figure 7.**
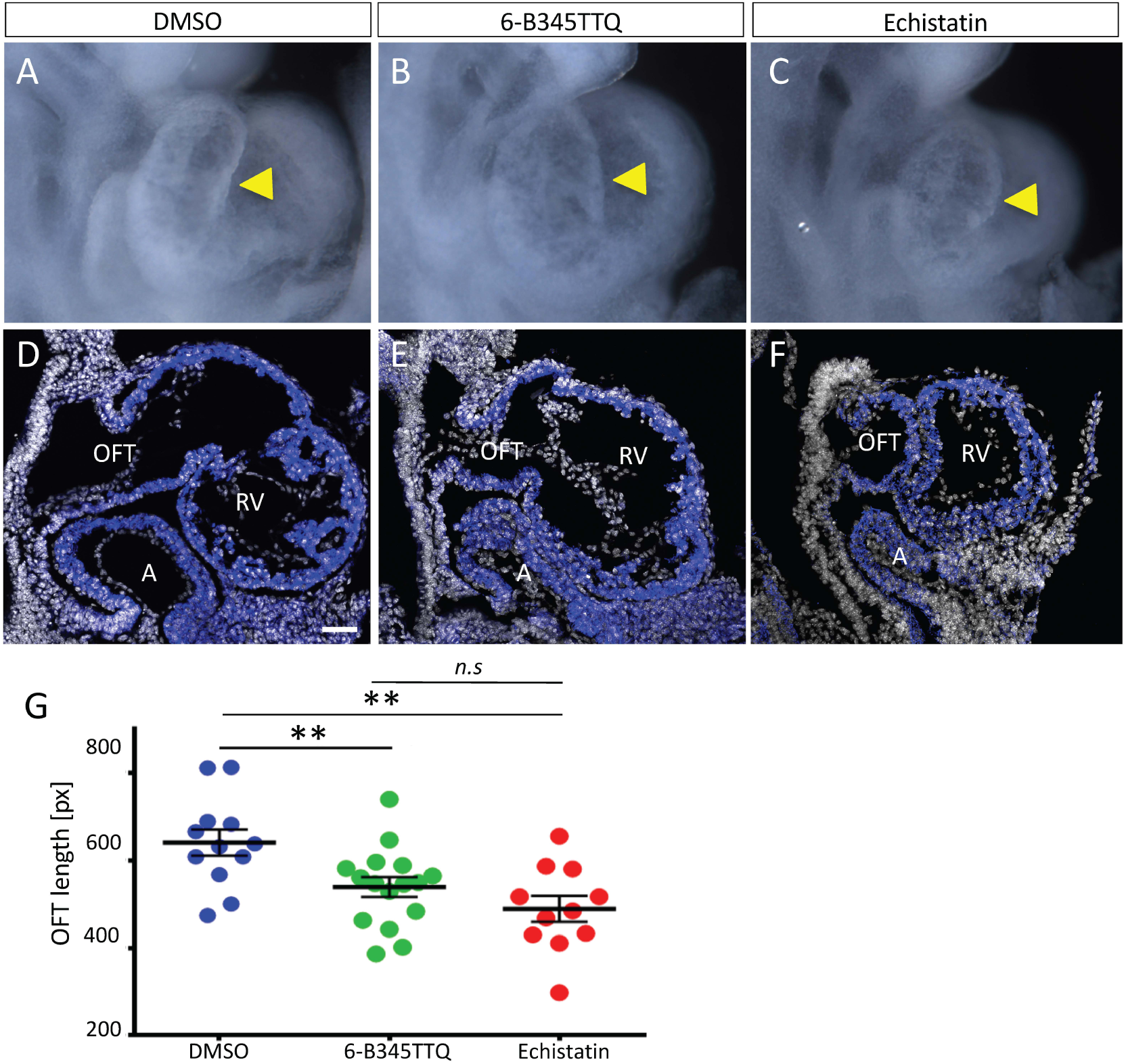
Integrin signalling is required for normal OFT development. (**A-C**) Right view of E9.5 embryos showing OFT (arroweads), cultured *ex-vivo* for 20h, control (DSMO) and treated embryos. (**D-F**) Sagittal section showing N-cadherin staining (blue) in heart tube and DPW in integrin inhibitor-treated compared with control embryos. Scale bars: A-F: 50 µm. (**G**) Measurement of OFT length of treated embryos (n=12 DMSO-treated embryos; n= 16 6-B345TTQ-treated, n=11 Echistatin-treated). P<0.01 (bilateral Mann-Whitney test). Error bars represent s.e.m. A, atrium; OFT, outflow tract; RV, right ventricle.

## DISCUSSION

*Tbx1* is required for cardiac progenitor cell contribution to the arterial and venous poles of the heart in mouse embryos, and is a major candidate gene for 22q11.2 deletion syndrome, which is often associated with congenital heart disease. In this work, we show that TBX1 is a regulator of the ECM-integrin-FA-intracellular signalling axis in SHF cells of the SpM of mouse embryos, and that loss of TBX1 has disruptive consequences on tissue architecture and heart development. The ECM-integrin-FA axis is central for cell migration, adhesion and polarity, and we show that in an embryo culture model, inhibition of integrin-FA interactions in a specific time window is associated with OFT dysmorphogenesis. Together our results indicate that abnormalities of ECM-integrin-FA signaling are likely to contribute to the OFT defects caused by loss of TBX1, providing new mechanistic insights into the origin of congenital heart defects.

*Tbx1* is expressed in SpM in both the epithelial layer and adjacent mesenchyme. Previous reports have hypothesized a process of intercalation as a means of mesenchymal cells to contribute to the epithelial layer (15-17). However, this model still needs to be confirmed. Given the continuity of gene expression of several SHF markers in the mesenchyme and epithelial layer, and the discontinuity of the basal membrane, it is possible that mesenchymal cells contribute to the epithelial layer through a process akin to condensation. This process is mediated by cadherin in a manner that is mechanistically analogous to integrin-mediated spreading on ECM (31). Live imaging will address the cell dynamics of eSHF expansion and putative contribution from the underlying mesenchyme.

We have characterized novel features of the epithelial properties of the SHF and how dysregulation causes defects at both the cellular and tissue-wide levels. In particular we found that FA proteins PXN and VCL are localized at the apical and basal sides of the DPW/eSHF. The basal localization is consistent with typical FA functions in the interaction between ECM-integrin-FA-cytoskeleton components. However, the apical localization and its extensive overlap with E-cadherin and beta-catenin are consistent with association with actin cytoskeleton at epithelial junctions or cell-cell junctions, similar to what has been described in other epithelia (23), (32-42). The finding that ITGB1 mostly accumulates at cell-cell contacts in the eSHF suggests the existence of a functional interaction between integrin- and cadherin-mediated cell adhesion complexes. This adhesion complex might represent an anchor point for mesenchymal cells in the eSHF either during initial mesenchymal condensation and/or during deployment of SHF cells to the cardiac poles.

At the apical/lateral region of the eSHF we also found a contractile protein, NMIIB/MYH10, a non-muscle myosin that maintains cortical tension and cell shape, and is required for heart development (19, 21, 43, 44), although its specific role in the SHF has not been determined. The consequences of NMIIB mislocalization caused by loss of TBX1 are still unclear, but recent data has shown that disruption of the actomyosin machinery using a ROCK inhibitor changes cells shape but does not seem to affect epithelial tension in the SHF, consistent with a role for actomyosin in preventing cell distortion in the SHF in response to epithelial tension rather than regulating tension itself (14).

Our *in vivo* mosaic data indicate that at least some of the TBX1 functions in the SHF are non-cell autonomous, consistent with previously reported chimera analyses showing that individual *Tbx1*^-/-^ cells in a WT background are able to contribute to the OFT (11). These findings suggest a critical contribution of the ECM to the pathogenesis of phenotypic abnormalities and predict that these abnormalities may arise only if there is a critical number of mutant SHF cells. The crucial role of ECM-cell interaction in OFT development is supported by the OFT shortening observed following our integrin-FA blocking experiments. While integrin signaling has been shown to be essential for heart development (45-49), our data specifically implicate the SHF as a target cell type of integrin signaling during this process and identify a critical time window during which integrin-PXN interaction is required for OFT extension.

The identification of transcriptional targets of TBX1 provides mechanistic insight as to how this transcription factor may drive the processes described here. Indeed, ChIP-seq and RNA-seq data indicate that focal adhesion (including the *Pxn* gene), ECM-receptor and MAPK signaling are among the top-scoring pathways affected by reduced dosage of the *Tbx1* gene (50). Several of these pathways have been implicated in heart development. For example, conditional deletion of the FAK-encoding gene in the *Nkx2-5* expression domain is sufficient to cause OFT defects in mouse embryos (51). However, their upstream regulators during cardiac development have not yet been fully determined and it is likely that TBX1 regulates the ECM-integrin-FA axis by hitting multiple targets.

Overall, our data indicate that TBX1 impacts on cell dynamics and regulates ECM-intracellular signaling at different levels. In particular, we found that in the eSHF, PXN, VCL, E-cadherin, F-actin and NMIIB are localized at the apical-lateral region of the monolayer of cells, possibly assembling an actomyosin apparatus that provides, through cortical signaling, cohesiveness among the epithelial-like cells. The localization of most of these proteins and the polarity of eSHF cells is severely altered in mutant embryos, thus compromising the cohesiveness of the epithelial cell layer and providing a mechanistical basis for the previously observed cell roundness and protrusion phenotype (13). In addition, these polarity and cohesiveness defects could also explain the loss of tension within the SHF (14) and the reduced contribution of SHF cells to the heart in these mutants. Our results thus reinforce and extend the emerging paradigm that regulation of the epithelial properties of cardiac progenitor cells in the SHF is a critical step in early heart morphogenesis. Moreover, they provide new insights into how SHF regulators such as TBX1 may impact on progenitor cells properties including proliferation and differentiation through the control of ECM-cell interactions and tissue architecture.

## EXPERIMENTAL PROCEDURES

### Mouse lines

The mouse lines *Tbx1*^*lacZ*/+^ (null allele, here referred to as *Tbx1*^-/+^) (52), *Tbx1*^*mcm*/+^ (null allele with Tamoxifen-inducible Cre knock in) (11), and *Tbx1*^*flox*/+^ (conditional floxed allele) (11) were maintained in a clean facility in a C57Bl/6N background. All mouse lines utilized here are available through public repositories: *Rosa*^*mT/mG*^, The Jackson Laboratory stock N: 007576, *Tbx1*^*mcm*^, EMMA repository EM:02404, *Tbx1*^*flox/flox*^, EMMA repository EM:02135, *Tbx1*^*Lacz*^, EMMA repository EM:02137. Genotyping was carried out according to instructions provided by the original reports. The developmental stage of embryos was evaluated by considering the morning of vaginal plug as embryonic (E) day 0.5, and by counting somites of embryos. Embryos were collected at E8.5 and E9.5. To induce nuclear translocation of the MerCreMer fusion protein encoded by the *Tbx1*^*mcm*^ allele, pregnant mice were intraperitoneally injected twice with Tamoxifen (75 mg/kg body weight) in the morning and afternoon of E7.5. Animal studies were carried out according to the animal protocol 257/2015-PR (licensed to the AB lab) reviewed by the Italian Istituto Superiore di Sanità and approved by the Italian Ministero della Salute, according to Italian regulations.

### Cell culture and transfections

Mouse C2C12 undifferentiated myoblast cells obtained from the ATCC (catalog CRL-1772) were cultured in Dulbecco’s modified Eagle’s medium (DMEM, Invitrogen) supplemented with 10% FBS and L-glutamine (300 *μ*g/ml), and were free of mycoplasma. Primary mouse embryonic fibroblasts (MEFs) were isolated from *Tbx1*^+/-^, *Tbx1*^-/-^, and wild-type embryos at E13.5. To this end, the internal organs, head, tail and limbs were removed. Cells were cultured in Dulbecco’s modified Eagle’s medium with 20% FBS and 1% NEAA, and used for a maximum of 3 passages. Cells were incubated at 37°C in 5% CO_2_. For siRNA transfection, cells were seeded at 1.2×10^5^ per well in six-well plates and transfected with a pool of Silencer Select Pre-Designed *Tbx1* SiRNA (Life Technology) (final concentration 50 nmol/L) in antibiotic-free medium using Lipofectamine RNAiMAX Reagent (Life Technology) according to the manufacturer’s instructions. 48hrs after siRNA transfection, cells were collected and processed for further analysis.

### Immunoblotting

Cells were harvested in lysis buffer (50 mmol/L Tris-HCl pH 7.6, 2 mmol/L EDTA, 150 mmol/L NaCl, 0.5% Triton X-100) supplemented with 1 mM phenylmethylsulphonyl fluoride and 1× complete mini EDTA-free protease inhibitor and PhosSTOP phosphatase inhibitor (Roche). Debris was removed by centrifugation at 10,000 *g* for 20 min at 4°C, and protein content was assessed by a Bradford protein assay. Nuclear proteins were extracted as described previously (23). Proteins were separated by SDS-PAGE and transferred to Immobilon-P PVDF membranes (Biorad). Membranes were subsequently incubated for 1 hour at room temperature in TBST buffer [125 mmol/L Tris-HCl (pH 8.0), 625 mmol/L NaCl, 0.1% Tween 20] containing 5% BSA and further incubated at 4°C for 16 hours with primary antibodies (listed in Table 1). Secondary HRP-conjugated mouse and rabbit (GE Healthcare) or rat (Dako) antibodies were used at a dilution of 1:5000. Membranes were developed using the enhanced chemiluminescence system (GE Healthcare). X-ray films were scanned and processed using ImageJ software for densitometric analysis.

**Table 1:** Primary antibodies for western blotting analysis.

|  |  |  |
| --- | --- | --- |
| Paxillin | Mouse | Thermo AHO0492 |
| Paxillin | Rabbit | Abcam ab32084 |
| Paxillin | Rabbit | Santa Cruz sc-5574 |
| E-cadherin | Mouse | BD #610182 |
| Vinculin | Mouse | Sigma #V9131 |
| Phospho-Pxn | Rabbit | Cell Signaling #2541 |
| FAK | Mouse | Santa Cruz #1688 |
| Phospho-FAK (Y925) | Goat | Santa Cruz #sc-11766 |
| Phospho-FAK (Y397) | Rabbit | Cell Signaling #8556 |
| Phospho-ERK (p44/42) | Rabbit | Cell Signaling #9101 |
| ERK | Rabbit | Cell Signaling #9102 |
| Integrin beta1<br>(cloneMB1.2) | Rat | Millipore MAB1997 |
| TBX1 | Rabbit | Abcam #ab18530 |
| Laminin B | Goat | Santa Cruz sc-6216 |
| GAPDH | Mouse | Abcam #125247 |
| Cofilin1 | Rabbit | Cell Signaling #3318 |
| Phospho-Cofilin1 | Rabbit | Cell Signaling #3311 |
| DDR1 | Rabbit | Santa Cruz #sc-532 |

### Migration assay

Transwell filters coated with 0.1% gelatin (diluted in PBS) with a PET membrane with 8 μm pores (BD Biosciences) were rehydrated for 2 hrs at 37°C in medium without supplements. Transfected cells were washed once in PBS, and 10^5^ cells were seeded into the upper chamber of the Transwells in serum-free medium containing 0.1% BSA. Medium containing 10% FBS or 0.1% BSA was added to the bottom chamber as a chemoattractant. Control wells without FBS were included to assess random migration. Cells were allowed to migrate for 4 hrs at 37°C, in 5% CO_2_. The cells on the bottom part of the membrane were fixed in methanol containing 0.1% Crystal Violet. Ten separate bright-field images were randomly acquired of each Transwell filter using a Leica DMI6000 microscope with a Plan APO 10×objective (Leica Microsystems). The cells in each image were counted and analyzed in comparison with control-transfected cells. Migration in the absence of FBS (random migration) was taken as 100% for each siRNA treatment, to control for variations in final cell number. For scratch wound assays cells were seeded 24 hrs after transfection in 24-well plates at confluency in growth medium; the confluent monolayer was wounded after 24 hrs by scraping with a pipette tip and then imaged by time-lapse microscopy. A phase-contrast image was acquired every 10 minutes for 18 hrs on a fully motorized Leica DMI6000 microscope with a Plan APO 10× objective. The area occupied by cells in scratch wounds was determined from time-lapse movie images taken at different time after wounding, using ImageJ analysis software. For polarity analysis, cells were plated at confluency on coverslip coated with Collagen I, wounded and fixed 4–6 hrs after wounding for anti-giantin immunostaining.

### Cell adhesion and spreading

Flat-bottom 96-well microtiter plates were coated with 10 μg/ml Collagen I (Sigma-Aldrich), 5 μg/ml Fibronectin (Roche), 100 μg/ml Matrigel (BD Biosciences) or 1% heat-denatured BSA in PBS (uncoated plastic) as a negative control, and incubated overnight at 4°C. Plates were then blocked for 1 hour at room temperature with 1% heat-denatured BSA in PBS. Cells were harvested using trypsin and allowed to recover for 1 hr. at 37°C, then washed three times in PBS. 10^5^ cells were plated in each coated well and incubated for 1 hr. at 37°C. Then, culture plates were washed with phosphate-buffered saline (PBS). Attached cells were fixed with 4% paraformaldehyde in PBS for 10 minutes. Cells were finally stained with 0.5% Crystal Violet in 20% methanol. The stain was eluted using 0.1 M sodium citrate in 50% ethanol, pH 4.2, and the absorbance at 595 nm was measured in a spectrophotometer. Alternatively, cells were fixed with 4% paraformaldehyde and counted under the microscope to corroborate the plate reader results. For cell spread area measurements, cells were allowed to spread on the indicated substrates for 20 min and fixed with paraformaldehyde without washing. Cells were then stained for F-actin, and area and perimeter quantified with ImageJ software.

### Flow Cytometry analysis

For analysis of surface-β1 integrin levels, cells were scraped into PBS, fixed in 4% PFA–PBS for 30 min on ice and fixed with 4% paraformaldehyde; then cells were blocked in 5% BSA–PBS for 30 min on ice. Cells were then incubated with 10 μg/ml anti-integrin β1 for 1 hr. on ice. Purified immunoglobulin was used as a negative control. The cells were then washed and incubated with a fluorescein isothiocyanate-labelled goat anti-rat IgG for 30 min at 4°C. Finally, the cells were washed and analyzed by flow cytometry using a FACSCanto (Becton Dickinson).

### Immunofluorescence and confocal microscopy

Cells were fixed with 4% paraformaldehyde, permeabilized with 0.2% Triton X-100 in PBS, and blocked with 1% BSA in PBS. After incubation with primary antibodies (listed in Table 2), cells were incubated (1hr, RT) with the appropriate secondary antibodies (Alexa Fluor 488, 594; 1:400; Molecular Probes) and, when indicated, with Alexa fluor phalloidin (wavelength 480 nm; Invitrogen) for F-actin visualization; DAPI was used for nuclear staining. Confocal images were acquired with an inverted confocal microscope (Nikon A1) using 63× (1.4 NA) objective lenses. Images were from cells fixed 48 hrs after transfection unless indicated differently.

**Table 2:**
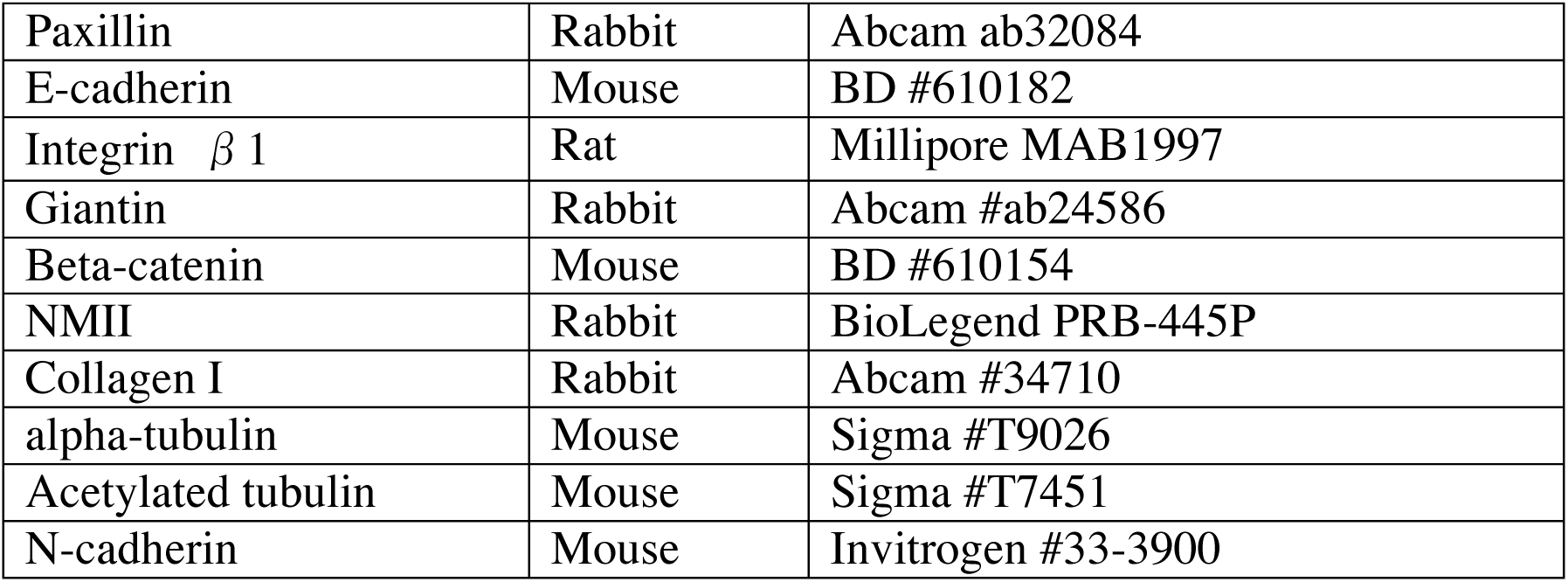
Primary antibodies for immunofluorescence analysis.

### FAs size analysis

A pipeline of Cell Profiler software (53) was used to quantify focal adhesions/complexes (number and size) from images of cells stained with anti-Paxillin, anti-phospho-Paxillin or anti-vinculin antibodies. Cells were either serum starved and stimulated to spread onto ECM-coated coverslips or grown in medium containing 10% FBS on glass coverslips. More than 100 FAs from two different experiments were analyzed.

### Histology and Immunofluorescence

E9.5 (23-25 somites) mouse embryos were fixed in 4% PFA and embedded in OCT. 10μm sagittal or transverse cryosections and subjected to immunofluorescence using primary antibodies listed in Table 2), and secondary antibodies Alexa fluor 488 (or 594 or 647) Goat anti-rabbit (or anti-mouse or anti-rat) IgG (H+L) (Life Technologies, 1:400). Alternatively, embryos were embedded in paraffin and then sectioned for immunostaining. Immunofluorescence experiments were imaged using a NIKON A1 confocal microscope or a Zeiss LSM 780 confocal microscope. We tested at least 3 embryos per antibody per genotype.

### Embryo culture

Embryo culture was performed as previously described (28). Embryos were cultured for 20 hours from E8.5 (7-11 somites). Control embryos were cultured with DMSO and treated embryos were cultured with either 50*μ*M 6-B345TTQ (B7438, Sigma-Aldrich), a specific inhibitor of paxillin-integrin*α*4 interaction or with 2.5*μ*g/ml of Echistatin (E1518, Sigma-Aldrich), a potent disintegrin.

### Measurements and quantifications of OFT

Right lateral views of whole embryos were used to measure OFT length using ImageJ. Number of embryos per condition: 12 DMSO-cultured, 16 6-B345TTQ-cultured and 11 Echistatin-cultured embryos. Data are presented as mean±s.e.m. P values were obtained using a bilateral Mann–Whitney test.

### RNA extraction, cDNA synthesis, and quantitative RT-PCR

RNA was extracted from C2C12 cells using TRI-Reagent (Ambion/Applied Biosystems) according to the manufacturer’s protocol. Extracted RNA was treated with DNA-free Kit (Ambion/Applied Biosystems). cDNA was synthesized from 1 μg total RNA (normalized via UV spectroscopy) using the High Capacity cDNA Reverse Transcription Kit, according to the manufacturer’s instructions (Applied Biosystems). Target cDNA levels were compared by Q-RT-PCR in 20-μl reactions containing 1× SYBR green (FastStart Universal SYBR Green Master (Rox), Roche), 0.2 μmol/L of each primer. Primers were as follows: for *Tbx1* amplification, forward primer 5’-CTGACCAATAACCTGCTGGATGA-3’ and reverse primer 5’-GGCTGATATCTGTGCATGGAGTT-3’; for *Pxn* amplification, forward primer 5’-GCTGCTGCTTCTGCTTCATC-3’ and reverse primer 5’-GTGGGTCCTCATTGGTCTGG-3’; for *Rpl13a* amplification, forward primer 5’-CCCTCCACCCTATGACAAGA-3’ and reverse primer 5’-CTGCCTGTTTCCGTAACCTC-3’. Results were normalized against *Rpl13a* and compared by relative expression and the delta-delta-cycle threshold method for fold change calculations with the StepOne v2.3 software (Applied Biosystems).

### Statistical analysis

Statistical significance was determined by a two-tailed paired Student’s t-test, unless otherwise specified. P-values <0.05 were considered as statistically significant. Error bars represent s.d.

### Data availability

All the data used in this work are shown in the Figures and in the Supplementary information. All other data supporting the findings of this study are available from the corresponding authors on reasonable request.

## AUTHOR CONTRIBUTIONS

D. A., A.A., C.C. and M.B. performed the experiments. D.A., C.C., K.R.G. and A.B planned and designed the experiments. D.A., C.C., K.R.G. and A.B. analysed the data. D.A., C.C., K.R.G. and A.B. jointly conceived and designed the study, and wrote the manuscript. All authors reviewed and approved the manuscript.

## ACKNOWLEDGEMENTS

We thank Maddy Parsons for critical reading of the manuscript, and Rosa Ferrentino for excellent technical assistance. We acknowledge the support of the Mouse Facility, Integrated Microscopy, and FACS Facility of IGB-CNR.

## SUPPLEMENTARY FIGURE LEGENDS

**Supplementary Figure 1. COL1 expression in SpM and DPW.**

(**A-B**) Transverse sections of E9.5 embryos showing the distribution of COLI (green) (arrows) at the SHF level in WT (A) and *Tbx1*^-/-^ embryos (B). (scale bar, 100μm). NT, neural tube; PE, pharyngeal endoderm; IFT, inflow tract.

**Supplementary Figure 2. COL1 and ITGB1 expression in the DPW/SpM.**

(**A-B**) Transverse sections of E9.5 embryos showing the distribution of COLI (green) and ITGB1 (red) at the SHF level in WT (A) and *Tbx1*^-/-^ embryos (B). (scale bar, 100μm). PE, pharyngeal endoderm; DPW, dorsal pericardial wall. C) sagittal section of a E9.5 WT embryo showing the distribution of ITGB1 and COLI. At basal side of eSHF cells is shown a partial colocalization between the two proteins (scale bar, 30 μm). DPW, dorsal pericardial wall; OFT, outflow tract; IFT, inflow tract; A, anterior; P, posterior; V, ventral; D, dorsal.

**Supplementary Figure 3. TBX1 is required to maintain actomyosin distribution and cell polarity in the eSHF.**

**(A-B)** Sagittal sections showing F-actin staining. In the WT (A) F-actin is localized at the cell cortex (arrows); in the *Tbx1*^-/-^ embryo (B) the staining is strongly reduced in the eSHF. Nuclei were stained with DAPI (scale bar, 10μm). **(C-D)** Confocal images of immunofluorescence of sagittal sections of E9.5 WT and *Tbx1*^-/-^ embryos stained with NMIIB antibody. NMIIB is apically distributed in WT embryos (arrows) while in the *Tbx1*^-/-^ embryo the protein surrounds the entire perimeter of the cells (arrows) (scale bar, 40μm). **(E-F)** Immunofluorescence images of sagittal sections of E9.5 WT and *Tbx1*^-/-^ embryos by using a Giantin antibody. (E) In the WT embryo, Giantin is oriented towards the pericardial cavity as its major mass was facing apically (parallel to the red dashed line) (arrow). (F) In the *Tbx1*^-/-^ embryo it is localized mostly laterally to the cells (arrow) (scale bar, 20μm). **(G)** Quantitative analysis of Giantin polarization data. Histograms show the mean ± s.d. of >100 cells in 3 embryos. Values are shown relative to control (\**p* < 0.05). SHF, second heart field; A, anterior; P, posterior; V, ventral; D, dorsal.

**Supplementary Figure 4. TBX1 regulates NMIIB and CDH1 distribution in the SHF in a cell non-autonomous manner.**

(**A-B**) Timed fate mapping of *Tbx1* in E9.5 *Tbx1*^*mcm*/+^; *R26R* embryos exposed to TM at E7.5, stained with NMIIB (green) and GFP (red) antibodies. GFP+ *Tbx1* homozygous cells (A-A’) and GFP+ *Tbx1* heterozygous cells (B-B’) show identical apical NMIIB localization (arrows) (scale bars, 20μm). The regions shown in A and in B are boxed in A’ and B’, respectively (scale bars, 10μm). Single channels of this picture are shown in Supplementary Fig. 4E. **(C-D)** Timed fate mapping of *Tbx1-*expressing cells in E9.5 *Tbx1*^*mcm*/+^; *R26R* embryos exposed to TM at E7.5, stained with GFP (green) and CDH1 (red) antibodies. GFP+ *Tbx1* homozygous cells (C) and GFP+ *Tbx1* heterozygous cells (D) show similar CDH1 staining (arrows) (scale bars, 20μm). Single channels of this picture are shown in Supplementary Fig. 4F. SHF, second heart field; OFT, outflow tract; A, anterior; P, posterior; V, ventral; D, dorsal.

**Supplementary Figure 5. Effects of *Tbx1* knockdown on C2C12 cell proliferation, polarity and actin dynamics.**

**(A)** Western blot showing reduced expression of TBX1 after 48 hrs. from siRNA transfection. **(B)** After 24h from siRNAs transfection, C2C12 cells were maintained in culture for 16hrs, 24hrs or 48hrs and then subjected to a proliferation assay (CyQuant). Values are expressed as relative to control NT. Values are the means ± s.d. of three experiments performed in triplicate (*p< 0.05). (**C**) The histogram shows quantification of Giantin polarization data: the Giantin signal was counted as oriented towards the wound (polarized cells) if its major mass was facing the wound edge (scale bar, 50μm). Results are means ± s.d. from three independent experiments (>300 cells/each). (**D-E**) Representative pictures of alpha-tubulin immunofluorescence in control (NT) or *Tbx1*^*KD*^ cell. Arrows and arrowheads indicate leading or trailing edge of the control cell, respectively (scale bar, 25μm). (**F**) Flow cytometry-based quantification of forward scatter (FSC) and side scatter (SSC) parameters (means ± s.d., n=3). (**G-H**) Confocal images of siRNAs-treated cells seeded on COLI and stained with anti-PXN and anti-ITGB1 integrin antibodies. Nuclei were stained with DAPI. Arrows indicate FAs (scale bar, 15μm). (**I-M**) Aberrant cell protrusions in *Tbx1*^*KD*^ cells seeded on plastic (white columns) or COLI (black columns): (I-L) photomicrographs or (M) quantification (scale bar, 50μm). Values are the mean ± s.d. of three experiments (* *p* < 0.05).

**Supplementary Figure 6. Actin protrusions, microtubule arrangements and ECM-dependent FA signalling are altered in *Tbx1*-depleted C2C12 cells.**

(**A**) Quantitative analysis of membrane lamellipodial protrusions in the migrating C2C12 cell sheet, measured by two different parameters: Lamellipodial density or number of cell protruding facing wound margin. Values are the means ± s.d. of two independent experiments (*p< 0.05). (**B-E**) Representative pictures of microtubules immunostained with alpha-tubulin (right panels) or anti-acetylated-tubulin (left panels). In all images PXN is shown in green. (scale bar, 15μm). (**F**) After 48hrs from siRNAs transfection, C2C12 cells were plated on COLI or FN and lysed after 20 min. Cell lysates were analysed by immunoblotting with Pospho-CFL1 antibody. The levels were normalised to total CFL1. GAPDH was used as a loading control. (**G**) After 48 hrs from siRNAs transfection, C2C12 cells were plated on COLI or FN and lysed after 20 min. Cell lysates were analysed by immunoblotting with antibodies to the indicated phosphorylated proteins. GAPDH was used as loading control. *Pxn*-depleted cells were used as positive controls of the experiment.

**Supplementary Figure 7. Integrin-FA pathway is critical for OFT elongation.**

(**A**) Scheme of embryo culture experiment. (**B**) Right view of E9.5 embryo, cultured *ex-vivo* for 20h; measurement of OFT length of embryo (yellow arroweads). (**C**) Examples of cultured embryos: whole mount photographs (upper panels). The table summarizes the number of embryos measured (DMSO refers to a control set) (lower panel).

**Supplementary Movie 1 and Movie2.**

3D reconstructions of PXN (green) and CDH1 (red) immunostaining in outer layer of SHF of a WT embryo (Nikon A1 confocal software).

## Non-standard abbreviations and Acronyms

CDH1: E-cadherin
COLI: collagen type I
E: Embryonic day
ECM: extracellular matrix
FA: focal adhesion
GFP: green fluorescent protein
IF: immunofluorescence
NMIIB: non-muscle myosin II isoform B (MYH10)
OFT: outflow tract
PXN: Paxillin
PCR: polymerase chain reaction
SHF: second heart field
TBX1: T-box transcription factor 1
VCL: vinculin
WT: wild-type

## REFERENCES

1. Meilhac SM, et al. (2003) A retrospective clonal analysis of the myocardium reveals two phases of clonal growth in the developing mouse heart. Development 130(16):3877–3889.

2. Buckingham M, Meilhac S, & Zaffran S (2005) Building the mammalian heart from two sources of myocardial cells. Nat Rev Genet 6(11):826–835.

3. Meilhac SM, Esner M, Kelly RG, Nicolas J-F, & Buckingham ME (2004) The Clonal Origin of Myocardial Cells in Different Regions of the Embryonic Mouse Heart. Dev Cell 6(5):685–698.

4. Zaffran S, Kelly RG, Meilhac SM, Buckingham ME, & Brown NA (2004) Right ventricular myocardium derives from the anterior heart field. Circ Res 95(3):261–268.

5. Vincent SD & Buckingham ME (2010) How to make a heart: the origin and regulation of cardiac progenitor cells. Curr Top Dev Biol 90:1-41.

6. Lescroart F, et al. (2014) Early lineage restriction in temporally distinct populations of Mesp1 progenitors during mammalian heart development. Nat Cell Biol 16(9):829– 840.

7. Devine WP, Wythe JD, George M, Koshiba-Takeuchi K, & Bruneau BG (2014) Early patterning and specification of cardiac progenitors in gastrulating mesoderm. Elife 3:e03848.

8. Diogo R, et al. (2015) A new heart for a new head in vertebrate cardiopharyngeal evolution. Nature 520(7548):466–473.

9. De Bono C, et al. (2018) T-box genes and retinoic acid signaling regulate the segregation of arterial and venous pole progenitor cells in the murine second heart field. Hum Mol Genet:ddy266–ddy266.

10. Cortes C, Francou A, De Bono C, & Kelly RG (2018) Epithelial Properties of the Second Heart Field. Circ Res 122(1):142–154.

11. Xu H, et al. (2004) Tbx1 has a dual role in the morphogenesis of the cardiac outflow tract. Development 131(13):3217–3227.

12. Rana MS, et al. (2014) Tbx1 coordinates addition of posterior second heart field progenitor cells to the arterial and venous poles of the heart. Circ Res 115(9):790–799.

13. Francou A, Saint-Michel E, Mesbah K, & Kelly RG (2014) TBX1 regulates epithelial polarity and dynamic basal filopodia in the second heart field. Development 141(22):4320–4331.

14. Francou A, De Bono C, & Kelly RG (2017) Epithelial tension in the second heart field promotes mouse heart tube elongation. Nat Commun 8:14770.

15. Sinha T, Wang B, Evans S, Wynshaw-Boris A, & Wang J (2012) Disheveled mediated planar cell polarity signaling is required in the second heart field lineage for outflow tract morphogenesis. Dev Biol 370(1):135–144.

16. Sinha T, et al. (2015) Loss of Wnt5a disrupts second heart field cell deployment and may contribute to OFT malformations in DiGeorge syndrome. Hum Mol Genet 24(6):1704–1716.

17. Li D, et al. (2016) Spatial regulation of cell cohesion by Wnt5a during second heart field progenitor deployment. Dev Biol 412(1):18–31.

18. Chen L, et al. (2012) Transcriptional control in cardiac progenitors: Tbx1 interacts with the BAF chromatin remodeling complex and regulates Wnt5a. PLoS Genet 8(3):e1002571.

19. Ma X & Adelstein RS (2012) In vivo studies on nonmuscle myosin II expression and function in heart development. Front Biosci (Landmark Ed) 17:545-555.

20. Marigo V, et al. (2004) Correlation between the clinical phenotype of MYH9-related disease and tissue distribution of class II nonmuscle myosin heavy chains. Genomics 83(6):1125–1133.

21. Phillips HM, Murdoch JN, Chaudhry B, Copp AJ, & Henderson DJ (2005) Vangl2 acts via RhoA signaling to regulate polarized cell movements during development of the proximal outflow tract. Circ Res 96(3):292–299.

22. Ridley AJ, et al. (2003) Cell migration: integrating signals from front to back. Science 302(5651):1704–1709.

23. Yano H, et al. (2004) Roles played by a subset of integrin signaling molecules in cadherin-based cell-cell adhesion. J Cell Biol 166(2):283–295.

24. Owen KA, et al. (2007) Regulation of lamellipodial persistence, adhesion turnover, and motility in macrophages by focal adhesion kinase. J Cell Biol 179(6):1275–1287.

25. Giancotti FG & Ruoslahti E (1999) Integrin signaling. Science 285(5430):1028–1032.

26. Hynes RO (2002) Integrins: bidirectional, allosteric signaling machines. Cell 110(6):673–687.

27. Bamburg JR (1999) Proteins of the ADF/cofilin family: essential regulators of actin dynamics. Annu Rev Cell Dev Biol 15:185-230.

28. Dominguez JN, Meilhac SM, Bland YS, Buckingham ME, & Brown NA (2012) Asymmetric fate of the posterior part of the second heart field results in unexpected left/right contributions to both poles of the heart. Circ Res 111(10):1323–1335.

29. Liu S, et al. (1999) Binding of paxillin to alpha4 integrins modifies integrin-dependent biological responses. Nature 402(6762):676–681.

30. Kummer C, Petrich BG, Rose DM, & Ginsberg MH (2010) A small molecule that inhibits the interaction of paxillin and alpha 4 integrin inhibits accumulation of mononuclear leukocytes at a site of inflammation. J Biol Chem 285(13):9462–9469.

31. Gumbiner BM (1996) Cell adhesion: the molecular basis of tissue architecture and morphogenesis. Cell 84(3):345–357.

32. Tornavaca O, et al. (2015) ZO-1 controls endothelial adherens junctions, cell-cell tension, angiogenesis, and barrier formation. J Cell Biol 208(6):821–838.

33. Crawford BD, Henry CA, Clason TA, Becker AL, & Hille MB (2003) Activity and distribution of paxillin, focal adhesion kinase, and cadherin indicate cooperative roles during zebrafish morphogenesis. Mol Biol Cell 14(8):3065–3081.

34. Bazellieres E, et al. (2015) Control of cell-cell forces and collective cell dynamics by the intercellular adhesome. Nat Cell Biol 17(4):409–420.

35. le Duc Q, et al. (2010) Vinculin potentiates E-cadherin mechanosensing and is recruited to actin-anchored sites within adherens junctions in a myosin II-dependent manner. J Cell Biol 189(7):1107–1115.

36. Yamada S, Pokutta S, Drees F, Weis WI, & Nelson WJ (2005) Deconstructing the cadherin-catenin-actin complex. Cell 123(5):889–901.

37. Barry AK, et al. (2014) alpha-catenin cytomechanics-role in cadherin-dependent adhesion and mechanotransduction. J Cell Sci 127(Pt 8):1779–1791.

38. Dorland YL & Huveneers S (2017) Cell-cell junctional mechanotransduction in endothelial remodeling. Cell Mol Life Sci 74(2):279–292.

39. Huveneers S, et al. (2012) Vinculin associates with endothelial VE-cadherin junctions to control force-dependent remodeling. J Cell Biol 196(5):641–652.

40. Kano Y, Katoh K, Masuda M, & Fujiwara K (1996) Macromolecular composition of stress fiber-plasma membrane attachment sites in endothelial cells in situ. Circ Res 79(5):1000–1006.

41. Playford MP, Vadali K, Cai X, Burridge K, & Schaller MD (2008) Focal adhesion kinase regulates cell-cell contact formation in epithelial cells via modulation of Rho. Exp Cell Res 314(17):3187–3197.

42. Watabe-Uchida M, et al. (1998) alpha-Catenin-vinculin interaction functions to organize the apical junctional complex in epithelial cells. J Cell Biol 142(3):847–857.

43. Ma X & Adelstein RS (2014) A point mutation in Myh10 causes major defects in heart development and body wall closure. Circ Cardiovasc Genet 7(3):257–265.

44. Tullio AN, et al. (1997) Nonmuscle myosin II-B is required for normal development of the mouse heart. Proc Natl Acad Sci U S A 94(23):12407–12412.

45. Mittal A, Pulina M, Hou S-Y, & Astrof S (2013) Fibronectin and integrin alpha 5 play requisite roles in cardiac morphogenesis. Dev Biol 381(1):10.1016/j.ydbio.2013.1006.1010.

46. Liang D, et al. (2014) Mesodermal expression of integrin alpha5beta1 regulates neural crest development and cardiovascular morphogenesis. Dev Biol 395(2):232–244.

47. Yang JT, Rayburn H, & Hynes RO (1995) Cell adhesion events mediated by alpha 4 integrins are essential in placental and cardiac development. Development 121(2):549– 560.

48. Sengbusch JK, He W, Pinco KA, & Yang JT (2002) Dual functions of α4β1 integrin in epicardial development. J Cell Biol 157(5):873.

49. Liu S, Rose DM, Han J, & Ginsberg MH (2000) α4 Integrins in Cardiovascular Development and Diseases. Trends in Cardiovascular Medicine 10(6):253–257.

50. Fulcoli FG, et al. (2016) Rebalancing gene haploinsufficiency in vivo by targeting chromatin. Nat Commun 7:11688.

51. Hakim ZS, et al. (2007) Conditional deletion of focal adhesion kinase leads to defects in ventricular septation and outflow tract alignment. Mol Cell Biol 27(15):5352–5364.

52. Lindsay EA, et al. (2001) Tbx1 haploinsufficieny in the DiGeorge syndrome region causes aortic arch defects in mice. Nature 410(6824):97–101.

53. Carpenter AE, et al. (2006) CellProfiler: image analysis software for identifying and quantifying cell phenotypes. Genome Biol 7(10):R100.

